# Whole Genome *De Novo* Variant Identification with FreeBayes and Neural Network Approaches

**DOI:** 10.1101/2020.03.24.994160

**Authors:** Felix Richter, Sarah U. Morton, Hongjian Qi, Alexander Kitaygorodsky, Jiayao Wang, Jason Homsy, Steven DePalma, Nihir Patel, Bruce D. Gelb, Jonathan G. Seidman, Christine E. Seidman, Yufeng Shen

## Abstract

**Motivation:** *De novo* variant (DNV) calling typically relies on heuristic filters intrinsic to specific platforms and variant calling algorithms. FreeBayes and neural network approaches have overcome this limitation for variant calling, and we implemented a similar approach for DNV identification.

**Results:** We developed a DNV calling framework that uses Genome Analysis Toolkit (GATK), FreeBayes and a neural network trained on Integrative Genomics Viewer pile-up plots (IGV-bot). We identified DNVs in 2,390 WGS trios and benchmarked results against heuristics based on GATK parameters. Results were validated *in silico* and with Sanger sequencing, with the latter showing true positive rates of 98.4% and 97.3% for SNVs and indels, respectively. Taken together we describe a scalable framework for DNV identification based on both FreeBayes and neural network methods.

**Availability:** Source code and documentation are available at https://github.com/ShenLab/igv-classifier and https://github.com/frichter/dnv_pipeline under the MIT license.

## Introduction

*De novo* genetic variations (DNVs) are highly relevant to human disease pathogenesis and can be identified by comparing DNA sequences between affected individuals and their parents. One standard method to identify DNVs is by filtering Genome Analysis Toolkit (GATK)^1^ variant calls to select sites with heterozygous child and homozygous parental genotypes. This filtering emphasizes sensitivity, so it is typically followed by a step that maximizes specificity: manual inspection of DNA pileup plots in Integrative Genomics Viewer (IGV) which is laborious for large variant sets.^2^ Sequence context variables identified with IGV can be abstracted to automated pipelines, but existing algorithms were optimized for exome sequencing.^3,4^

Determining the sequence context most salient to DNV identification is a challenge. This challenge has been addressed in non-DNV filtering with Bayesian and neural network machine learning approaches. Bayesian approaches to variant calling include those employed by GATK and FreeBayes (FB).^1,5^ FB differs from GATK by utilizing the literal reads mapped to a region rather than the alignment of reads, minimizing false insertion/deletion (indel) variant calls.

An orthogonal approach to variant identification utilizes neural networks.^6–8^ A neural network is a machine learning framework that captures complex feature dependencies for prediction and inference. Previous variant callers used curated features or images of DNA pile-up plots as inputs for neural network variant callers.

Overall, neural network methods and FreeBayes have been shown to account for sequence context across diverse platforms while maintaining higher variant calling accuracies than heuristic approaches, but these have not been reprised as a general framework for DNV identification. We improved on DNV identification by GATK from WGS data by using the intersection of calls with FreeBayes and a machine learning approach, termed IGV-bot. IGV-bot applies neural network learning to child/parent DNA pile-up plots from IGV. We demonstrate scalability by identifying high-quality candidate DNVs in 2,390 trios across four cohorts.

## Methods

### Candidate Variant Identification

We first identified potential DNVs by selecting GATK VQSR PASS variants (*i*.*e*., variants classified as true with an adaptive error model based on known true sites and artifacts) that were present in probands and absent from both parents. For the initial cohort (PCGC1, N=349 trios), GATK candidate variants were sub-divided into four evidence tiers using heuristics, allowing for benchmarking and sanity checks. Tier 1 variant heuristic filters were: (1) rare (AF ≤ 10^−4^ across all samples in 1000 Genomes, gnomAD exomes and gnomAD genomes); (2) 10-65 reads total, 7 alternate allele reads, and a 30-70% alternate allele ratio in the proband; (3) a minimum depth of 10 reads total and alternate allele ratio (AAR) < 5% in parents; (4) minimum genotype (GT) quality score of 60 in probands and 30 in parents; and (5) AC=1 across cohort. Tier 2 variants were broadened to those with minimum GT score 20, 20-80% AAR in the proband and maximum 10% AAR in parents. Tier 3 variants were extended to have at least 7 total reads, and a minimum alternative allele read count of 5 in the proband. In addition to these three tiers of evidence, separate heuristics were applied to optimize sensitivity for identifying candidate DNVs, labeled as “Alternative Tier”. The Alternative Tier parameters were GATK PASS, heterozygous ratio (AB) set to 0.2-0.8 in the proband, homozygous ratio (AB) less than 0.01 in both parents, depth (DP) 7-120, Joint Genotyping allele count (AC) = 1 across all trios, Genotype Quality (GQ) > 60 (proband), GQ > 30 (parents), Alternate Allele Depth (AAD) > 7 in the proband, and AAD < 3 in each parent. For the remaining three cohorts (N_PCGC2_=413, N_SSC1_=518, N_SSC2_=1,110) we employed the most lenient filters.

### FreeBayes Variant Calling in Candidate Variant Regions

Four hundred base pair regions centered on GATK candidate DNVs were submitted to FreeBayes for variant identification. DNVs jointly identified by GATK and FreeBayes were further filtered to remove variants that occurred in repetitive or low complexity regions. DNVs were filtered out if they occurred in repetitive sequences (homopolymers of at least 5 bases; 3 repeats of a dinucleotide or trinucleotide sequence), with simple repeat or low complexity regions as annotated by RepeatMasker, or that had close proximity to indel variants in the same individual (within 5 bps or overlapping). *In silico* visualization of 988 candidate DNVs was used to assess the efficacy of FreeBayes and sequence filters on false positive DNV calling rates.

### Convolutional neural network (CNN) data pre-processing, architecture, and model training

Putative DNVs were plotted with IGV, which served as inputs for IGV-bot. These red-green-blue pileup images of candidate variants capture and spatially organize reference and alternate allele reads. The images are used to classify variants as true or false as well as the type of variant (SNV, insertion, deletion, or complex). They are organized as child, mother, and father from top to bottom, and show the DNV locus plus 20 flanking base pairs both upstream and downstream (**Figures 1 and 2**). The images were further pre-processed by removing borders and metadata and keeping four flanking bases for every locus (two upstream, two downstream).

Following image generation, a training set was generated through manual curation/classification into one of six groups: fail (n=3,892), SNV (n=6,031), insertion (2,494), deletion (n=4,731), complex (n=802), uncertain (n=171), or inherited (n=916). These images were converted to a NumPy array and normalized to [0,1] by dividing this array by 255. After DNV pre-processing, the CNN architecture was specified. The CNN architecture was a standard architecture for the MNIST dataset, reprised here to categorize IGV plots into one of six classes of variants specified above. The CNN comprised 6 hidden layers, each with 32 convolutional filters, a kernel size of 3, a maximum pooling area size of 2, and a rectified linear unit activation function. Pooling was done after the 2^nd^, 4^th^, and 6^th^ layers, with a 0.25 dropout applied after each pooling to minimize overfitting. The final model compilation consisted of flattening to a 1-dimensional array, a 0.5 dropout, and a final soft-max activation function to determine the probability of every class. The MNIST results were used to initialize weights for every node.Having generated a model architecture, the model was trained independently five times (see **Figures 1, 2** for example training data). The final model was the training attempt with the lowest cross-validation cost function value. Every attempt consisted of training the model using two epochs and an 80:20 cross-validation split.

## Results

Among 349 trios, GATK identified 44,558 candidate DNVs among the 4 tiers (**Table 1**). This represented 128 DNVs/trio, including 91 single nucleotide variants (SNVs) and 37 insertion/deletions (indels). FreeBayes calling of candidate DNVs within the 400 bp region flanking GATK candidate DNVs followed by short repetitive element filtering kept 61% of variants (79% of SNVs and 16% of indels), resulting in 78 DNVs/trio. IGV-bot filtering kept 88% of the post-FreeBayes variants (89% SNVs, 77% indels). Consistent with both the FreeBayes and IGV-bot procedures capturing high quality DNVs, 82% of DNVs with the highest tier of evidence (*i*.*e*., GATK Tier 1) were kept, compared to only 2% for the lowest tier. The majority of candidate DNVs removed with FreeBayes and IGV-bot were indels, in contrast to the relatively modest changes in number of SNVs, as evidenced with both the DNVs/trio and total number of DNVs. In total, 53% of GATK DNVs were retained. These results were consistent in three other cohorts (N=2041 trios), with 56% of GATK DNVs retained after applying both FreeBayes and IGV-bot.

**Table 1.** DNV totals.

|  |  | **GATK dnvs** |  |  | **GATK-FB dnvs** |  |  | **GATK-FB-NN** |  |  |
| --- | --- | --- | --- | --- | --- | --- | --- | --- | --- | --- |
|  |  | **Total** | **SNV** | **Indel** | **Total** | **SNV** | **Indel** | **Total** | **SNV** | **Indel** |
| **PCGC1 (N=349)**<br>**Evidence Tier** | **Tier 1** | 26034 | 22333 | 3701 | 23199 | 21467 | 1732 | 21406 | 19942 | 1464 |
|  | **Tier 2** | 7918 | 3809 | 4109 | 2640 | 2403 | 237 | 1714 | 1592 | 122 |
|  | **Tier 3** | 5224 | 1163 | 4061 | 302 | 263 | 39 | 96 | 92 | 4 |
|  | **Alt. Tier** | 5772 | 4622 | 1150 | 1089 | 973 | 116 | 613 | 562 | 51 |
|  | **Total** | 44948 | 31927 | 13021 | 27230 | 25106 | 2124 | 23829 | 22188 | 1641 |
| **PCGC1**<br>**DNVs/trio** | **Mean** | 128.8 | 91.5 | 37.3 | 78.2 | 72.1 | 6.1 | 68.5 | 63.8 | 4.7 |
|  | **Max** | 258 | 164 | 170 | 133 | 125 | 17 | 119 | 114 | 14 |
|  | **Min** | 73 | 45 | 12 | 36 | 33 | 0 | 28 | 27 | 0 |
| **PCGC2 (N=413)** | **Total** | 73193 | 38672 | 34521 | 35530 | 32619 | 2911 | 33902 | 31302 | 2600 |
| **SSC1 (N=518)** | **Total** | 79380 | 54581 | 24799 | 42696 | 39401 | 3295 | 36011 | 33627 | 2384 |
| **SSC2 (N=1110)** | **Total** | 122764 | 96906 | 25858 | 86617 | 81418 | 5199 | 84027 | 79176 | 4851 |

*In silico* confirmation of the GATK, FreeBayes (GATK-FB), and IGV-bot (GATK-FB-NN) candidate DNVs was used to assess the relative specificity of the pipelines. Visualization of 505 candidate DNVs that were jointly called by GATK and FreeBayes (GATK-FB) confirmed presence of the *de novo* variant in 502 cases (99.8%) (**Table 2**). IGV visualization of 483 variants unique to the GATK candidate *de novo* variant set did not confirm *de novo* variants in 462 (95.7%), supporting the efficacy of the joint calling approach. Of the 21 GATK candidate DNVs that were not identified by FreeBayes but did confirm by IGV, 6 were not called by FreeBayes, 6 were in simple repeat regions, 5 were in regions of low complexity, and 2 each had a nearby indel or were low quality. Visualization of subsets of the 483 variants removed by specific filters agreed with 96/97 removed due to overlapping indels (98.7%), 11/21 removed due to nearby indel (53%), 13/16 removed due to dinucleotide repeat sequences, and 10/16 removed due to homopolymer sequences (62.5%). IGV-bot was trained on DNVs from PCGC1, so independent *in silico* confirmation was performed using 580 DNVs from PCGC2. Most variants true based on GATK-FB were kept with IGV-bot (95.4%). Among variants IGV-bot kept, 90% were true with *in silico* confirmation. In contrast, *in silico* confirmation agreed with 61% of variants removed by IGV-bot. These results are consistent with a high specificity (86%) and medium sensitivity (70%) on the already high-quality candidate DNVs identified with GATK and FreeBayes.

**Table 2.** Validation *in silico*.

**FreeBayes results for variants kept with GATK**
| *In Silico* Visualization Result | FB<br>Keep | All FB<br>removed |
| --- | --- | --- |
| True DNV | 502 | 21 |
| False DNV | 3 | 462 |
| Total | 505 | 483 |
| Proportion of variants kept with GATK | 0.606 | 0.394 |

**Subsets of FB remove variants**
| *In Silico* Visualization<br>Result | Overlapping Indel<br>Filter | Nearby Indel<br>Filter | Homopolymer | Dinucleotide<br>Repeat |
| --- | --- | --- | --- | --- |
| True DNV | 1 | 10 | 6 | 3 |
| False DNV | 96 | 11 | 10 | 13 |
| Total | 97 | 21 | 16 | 16 |

**IGV-bot results for variants kept with GATK and FreeBayes**
| *In Silico* Visualization Result | IGV-bot<br>keep | IGV-bot<br>remove |
| --- | --- | --- |
| True DNV | 264 | 112 |
| False DNV | 28 | 176 |
| Total | 292 | 288 |
| Proportion of variants kept with GATK/FB | 0.954 | 0.046 |

PCR validation was performed for 399 candidate DNVs (**Table 3**); Sanger sequencing was successful for 390. The true positive rate was lowest for GATK (95.4%), intermediate for GATK-FB (96.9%) and highest for GATK-FB-NN (98.1%). Notably, the improvement in true positive rate was primarily driven by improvements in indel calling, which increased from 87.5% to 97.3%, while staying similar for SNVs, with all methods having true positive rates of 98-99%.

**Table 3.** PCR validation results.

|  |  | **Total** | **SNV** | **Indel** |
| --- | --- | --- | --- | --- |
| Sanger results | Submitted | 399 | 289 | 110 |
|  | Sanger sequencing not successful<br>(no primer or PCR did not work) | 9 | 3 | 6 |
| GATK results | GATK *De Novo* | 390 | 286 | 104 |
|  | Sanger *De Novo* | 372 | 281 | 91 |
|  | **True Positive rate (%)** | **95.4** | **98.3** | **87.5** |
| GATK-FB | GATK-FB *De Novo* | 354 | 265 | 89 |
|  | Sanger *De Novo* | 343 | 261 | 82 |
|  | **True Positive rate (%)** | **96.9** | **98.5** | **92.1** |
| GATK-FB-NN | GATK-FB-NN *De Novo* | 318 | 244 | 74 |
|  | Sanger *De Novo* | 312 | 240 | 72 |
|  | **True Positive rate (%)** | **98.1** | **98.4** | **97.3** |

## Conclusion

Here we present a framework for DNV identification from WGS data using FreeBayes followed by a convolutional neural network. The results compared well with gold-standards of *in silico* filtering and PCR, comparing favorably to filtering using heuristics alone. Since these methods do not rely on parameters intrinsic to Illumina, they can be rapidly generalized to other sequencing platforms.

---

**Supplemental Figure 1.**
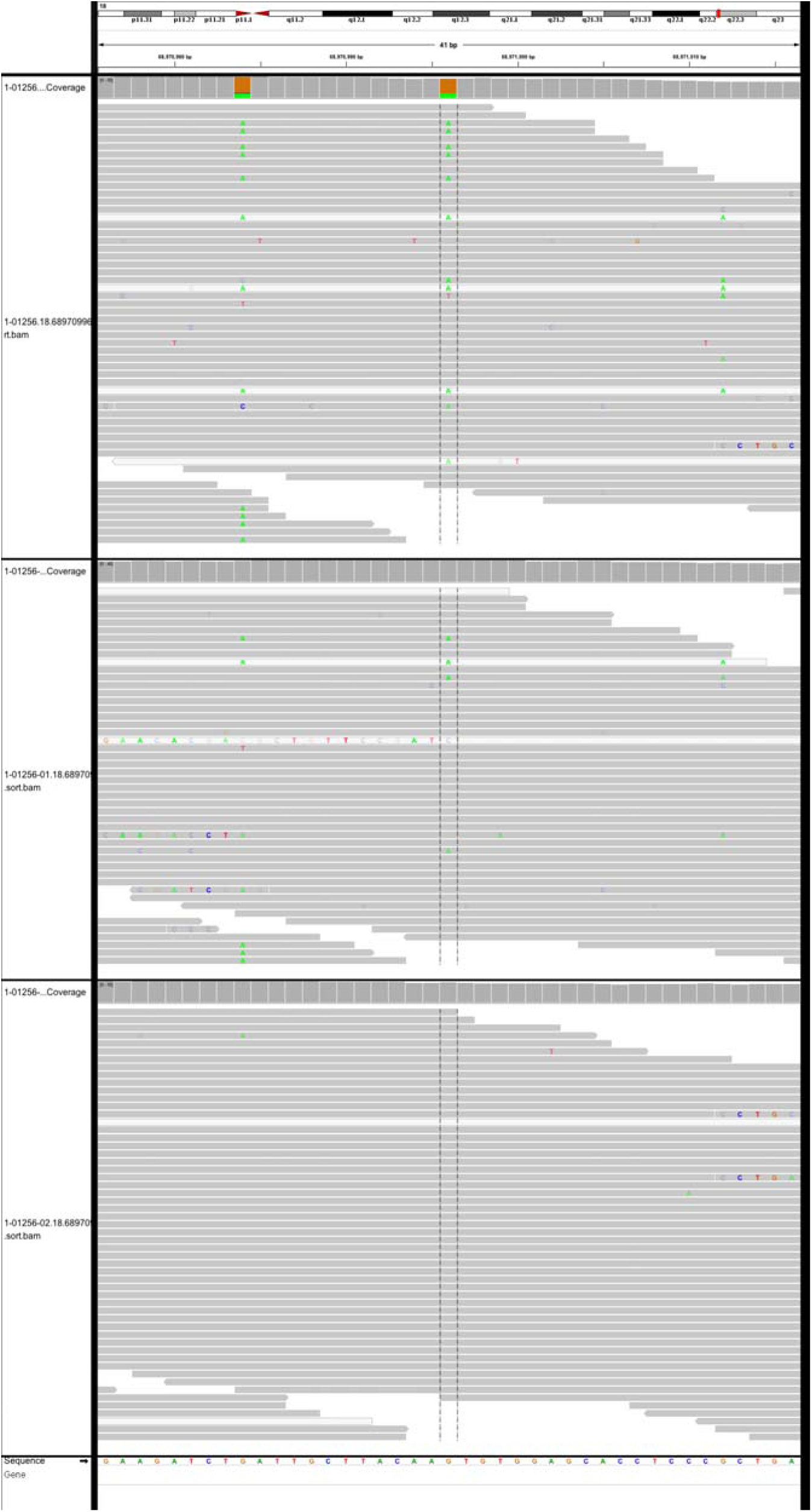
False variant. Example IGV pile-up of a false variant used to train the neural network, IGV-bot.

**Supplemental Figure 2.**
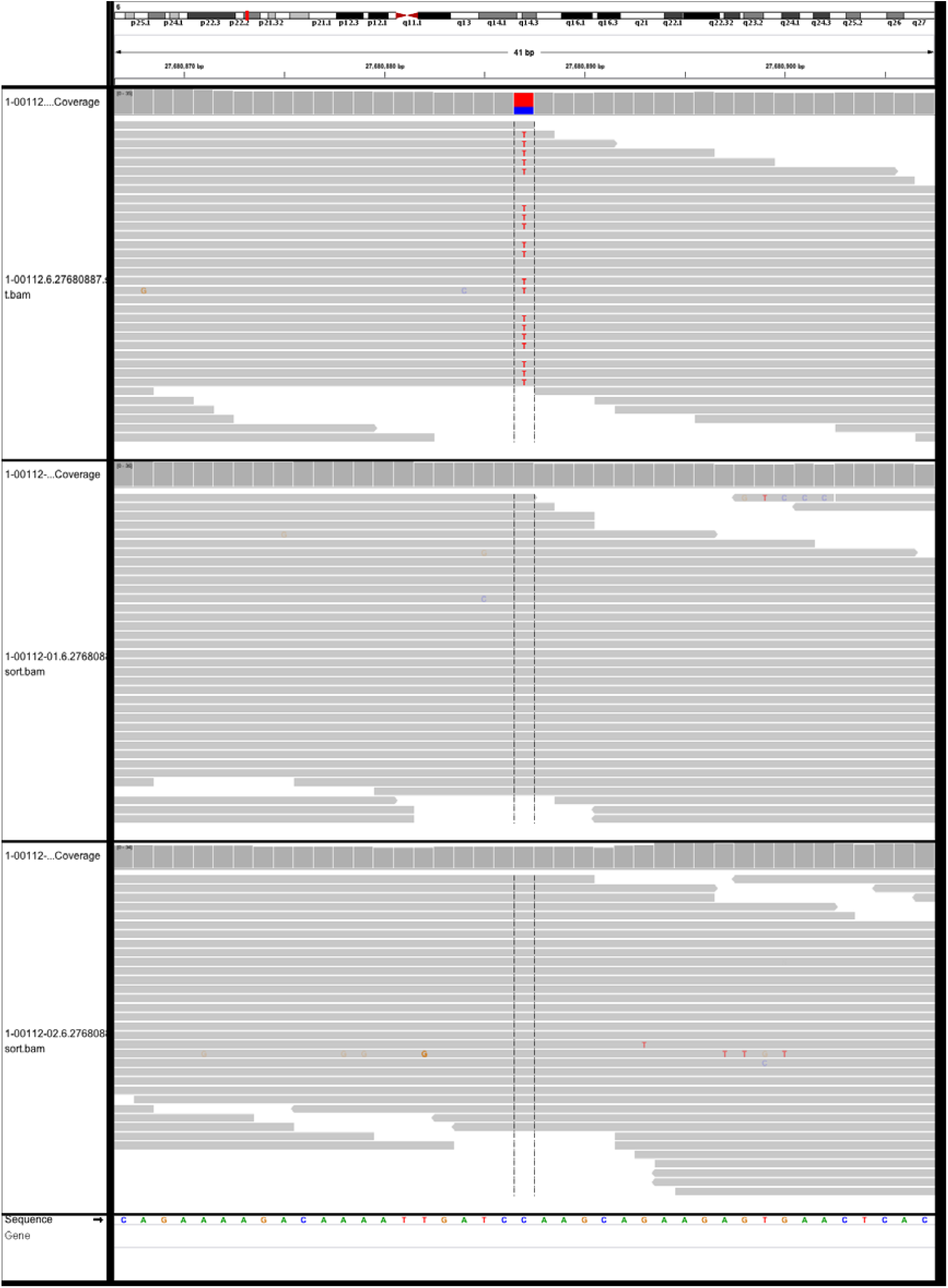
True SNV. Example IGV pile-up plot of a true single nucleotide variant used to train the neural network.

